# Vps501, a novel vacuolar SNX-BAR protein cooperates with the SEA complex to induce autophagy

**DOI:** 10.1101/2021.05.06.441257

**Authors:** Shreya Goyal, Verónica A. Segarra, Nitika, Aaron M. Stetcher, Andrew W. Truman, Adam M. Reitzel, Richard J. Chi

## Abstract

The sorting nexins (SNX), constitute a diverse family of molecules that play varied roles in membrane trafficking, cell signaling, membrane remodeling, organelle motility and autophagy. In particular, the SNX-BAR proteins, a SNX subfamily characterized by a C-terminal dimeric Bin/Amphiphysin/Rvs (BAR) lipid curvature domain and a conserved Phox-homology domain, are of great interest. In budding yeast, many SNX-BARs proteins have well-characterized endo-vacuolar trafficking roles. Phylogenetic analyses allowed us to identify an additional SNX-BAR protein, Vps501, with a novel endo-vacuolar role. We report that Vps501 uniquely localizes to the vacuolar membrane and works with the SEA complex to regulate autophagy. Furthermore, we found cells displayed a severe deficiency in starvation-induced/nonselective autophagy only when SEA complex subunits are ablated in combination with Vps501, indicating a cooperative role with the SEA complex during autophagy. Additionally, we found the SEA complex becomes destabilized in *vps501*Δ*sea1*Δ cells, which resulted in aberrant TORC1 hyperactivity and misregulation of autophagy induction.

## Introduction

The sorting nexin (SNX) family is an evolutionarily conserved class of cellular trafficking proteins that are most well-known for their ability to bind phospholipids to catalyze endosomal sorting reactions and other membrane trafficking pathways in the cell (Chi et al., 2015; Gallon and Cullen, 2015). SNX proteins are structurally characterized by an evolutionarily conserved region known as the Phox (PX) homology domain, which allows them to recognize the lipid composition of the endosome, most notably phosphatidylinositol-3-phosphate (PI3P) (Teasdale and Collins, 2012). While all PX domains have similar core folds, consisting of three antiparallel β-strands and three α-helices that extend a canonical PI3P binding motif ψPxxPxK (ψ=large aliphatic amino acid), outside this region, they have relatively low sequence homology. A recent analysis of over 39 PX domains identified a secondary His/Tyr-containing binding motif present in a subset of PX domain proteins which may explain the lipid promiscuity of some sorting nexins (Chandra et al., 2019). However, the range of different phosphoinositides that associate with PX domains, and the significance of these interactions for cellular localization are still unclear. Moreover, SNX proteins are divided into subfamilies according to the presence of other characteristic domains such as a Bin-Amphiphysin-Rvs (BAR) domain (van Weering et al., 2010). BAR domains allow members of the SNX-BAR subfamily to bind high positive curvature structures, driving the formation of endosomal tubules, while also conferring the ability to form sorting complexes that facilitate cargo selection (Chi et al., 2014; van Weering et al., 2012; van Weering et al., 2010). The *Saccharomyces cerevisiae* genome encodes seven annotated SNX-BAR proteins, while the human genome encodes twelve (Anton et al., 2020; Teasdale and Collins, 2012; van Weering et al., 2010). In addition to working with complexes such as retromer to mediate retrograde and recycling trafficking of cargos at the tubular endosomal network (TEN) (Burd and Cullen, 2014; Chi et al., 2014; Teasdale and Collins, 2012; van Weering et al., 2012; van Weering et al., 2010), SNX-BAR proteins contribute to other important conserved cellular processes such as macroautophagy (herein referred to as autophagy) and selective autophagy (Ma et al., 2017; Ma et al., 2018; Nemec et al., 2017).

Autophagy is a stress response in which eukaryotic cells recycle damaged or unneeded components by sequestering them in double-bilayered compartments called autophagosomes. Once made, autophagosomes deliver their contents for breakdown by docking and fusing with the cell’s degradative organelle, the lysosome in animal cells or the vacuole in plant and yeast cells. These key steps and the core autophagy-related (Atg) proteins that mediate and regulate them are evolutionarily conserved across all autophagy pathways, including starvation-induced bulk autophagy and cargo-selective autophagy pathways (Goyal et al., 2020; Nakatogawa, 2020; Su et al., 2015; Yang and Klionsky, 2010). While vacuoles and lysosomes can serve as storage and/or recycling depots for cells, their delimiting membranes host critical signaling events for autophagy induction such as the inactivation of *target of rapamycin* (TOR). Multiprotein complexes such as TORC1 in yeast are tethered to the vacuole membrane and function by integrating signals from many intracellular and extracellular cues from a variety of kinases, GTPases and their effectors (Binda et al., 2009). Recently, an upstream regulator of the TORC1, the yeast SEA complex (GATOR complex in humans), was identified and shown to be part of this web of GTPase effectors (Algret et al., 2014; Panchaud et al., 2013). The SEA complex is a conserved eight protein complex (Sea1, Sea2, Sea3, Sea4, Seh1, Sec13, Npr2, Npr3) made up of proteins with structural characteristics similar to the membrane coating complexes such as the nuclear pore complex, the COPII vesicle coating complex and HOPS/CORVET tethering complexes (Dokudovskaya and Rout, 2015). The SEA complex is also dynamically associated to the vacuole membrane; however, its complete function is not well understood.

Substantial effort has gone into understanding the membrane trafficking events required to form autophagosomes and the contributions of SNX-BAR proteins to the autophagy pathway. In fact, SNX-BAR proteins have been shown to mediate an emerging number of autophagy-related processes. For example, the Snx4-Snx41 SNX-BAR heterodimer mediates the retrograde endosome-to-Golgi transport of the autophagy-related protein Atg27, an integral membrane protein that when deleted leads to decreased autophagosome number and autophagic flux in budding yeast (Ma et al., 2017; Yen et al., 2007). Moreover, Snx4-mediated retrograde trafficking of proteins and lipids helps the cell maintain the phosphatidylserine (PS) and phosphatidylethanolamine (PE) homeostasis, which is required to allow autophagosome fusion with the vacuole, one of the final stages of autophagy (Ma et al., 2018). As we come to understand the full range of SNX-BAR protein functions, it is becoming clear that this group of proteins has a critical role in the network of cellular players that collectively regulate autophagy.

In this study, we report the identification of a novel yeast SNX-BAR protein, encoded by yeast open reading frame (ORF) YKR078W, that contributes to autophagy induction alongside the SEA complex. Our findings bring us to a more complete understanding of SNX-BAR family of proteins and the important role they play in autophagy regulation.

## Results

### YKR078W/VPS501 and VPS5 are phylogenetically related but functionally distinct

Phylogenetic analysis of the SNX-BAR sorting nexins allowed us to identify ORF YKR078W from *Saccharomyces cerevisiae* as a SNX-BAR candidate that forms a well-supported clade (bootstrap = 99) with Vps5 proteins from *S. cerevisiae* and other closely related species from the family Saccharomycetaceae (Figure 1A, Supplemental Figure 1). As previously shown by Koumandou et al. (2011), fungal Vps5 proteins are most closely related to SNX1/2 proteins from animals and choanoflagellates (Koumandou et al., 2011). The simplest explanation for the existence of two Vps5-like proteins in *S. cerevisiae* is a recent gene duplication event. To assess whether or not there is functional overlap between the Vps5 and YKR078W proteins, we used *YKR078WΔ* yeast cells to examine the localization of the vacuolar hydrolase receptor Vps10. This receptor is trafficked from the prevacuolar endosome to Golgi by the Vps5-dependent retromer-SNX-BAR complex and is mistrafficked to the vacuole in *vps5Δ* cells (Horazdovsky et al., 1997; Nothwehr and Hindes, 1997; Seaman et al., 1997). We hypothesized that, if there is functional overlap between the Vps5 and YKR078W proteins, Vps10 mislocalization in *YKR078WΔ* would phenocopy the *vps5Δ* mutant. However, deletion of YKR078W did not affect Vps10 localization (Figure 1B), indicating that while Vps5 and YKR078W are phylogenetically related, they are functionally different. In light of these findings, we gave YKR078W the distinct, but Vps5-related name of Vps501. We will use the Vps501 annotation throughout the remainder of this paper.

**Figure 1.**
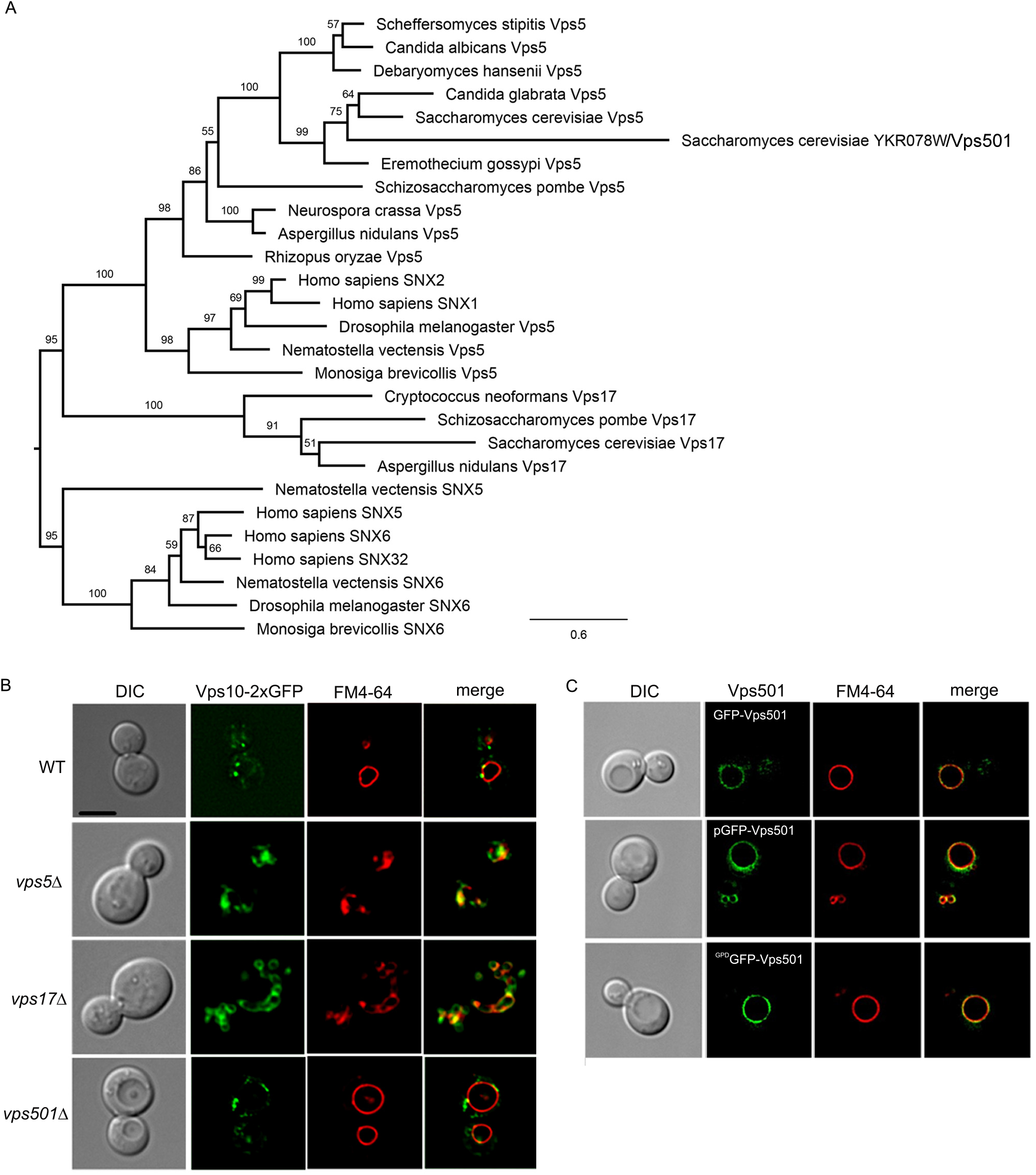
YKR078W/Vps501 is a paralog of Vps5 and resides on the vacuole membrane. (A) Phylogenetic analysis of Vps5-like SNX-BAR proteins in opisthokonts, animal, choanoflagellate and other family Saccharomycetaceae indicates the presence of a Vps5-like protein (YKR078W) in *Saccharomyces cerevisiae*, refereed here as Vps501. (B) Micrographs of Vps10-2XGFP in wildtype and indicated mutant cells. Vps501 does not have a role in Vps10 trafficking, despite the phylogenetic similarities with retromer SNX-BARs. (C) Vps501 localizes to the vacuole membrane as a N-terminal GFP fusion protein. GFP-Vps501 expression is shown as locus integrations using native promoter (top), GPD promoter (bottom) or ectopically expressed as a 2-micron plasmid (middle). C-terminal fusions were found to be non-functional, not shown. Vacuolar membranes are shown using FM4-64 dye. The scale bar indicates 5 μm. See also supplemental figure 1.

### N-terminally tagged Vps501 localizes to the vacuolar membrane

While Vps501 shares common evolutionary ancestors and lineage with Vps5, its function within the cell has so far remained uncharacterized. Multiple genome-wide localization screens using C-terminally fused GFP failed to detect expression and/or failed to localize Vps501 to any intracellular compartment (Huh et al., 2003). Interestingly, we have discovered that Vps501 appears to be nonfunctional as a C-terminal GFP fusion (data not shown). However, a fully integrated N-terminal GFP fusion appeared to preserve the functionality and localization of Vps501. Instead of an endosomal localization, which might be expected based on its homology to Vps5, we detect the protein predominantly on the limiting membrane of the vacuole, and in a few intracellular puncta (Figure 1C, upper panels). This localization pattern remains consistent when Vps501 is chromosomally tagged under the control of either its endogenous promoter (Figure 1C, upper panels) or a constitutive TDH3/GPD promoter (Figure 1C, lower panels), and when expressed from an extrachromosomal plasmid (Figure 1C, middle, panels).

### Identification of Vps501 interactors

To gain insights into Vps501 function, we used a co-immunoprecipitation mass spectrometry approach to identify its interactors. We purified GFP-Vps501 complexes and characterized the Vps501 interactome using mass spectrometry (Figure 2A). The list of Vps501 interacting proteins was enriched for proteins involved in amino acid/carbohydrate metabolism, protein translation, protein folding, and endocytosis (Supplemental Table 1). Our analysis of strong interactors of Vps501 identified subunits of the evolutionarily conserved eight-protein SEA complex (Sea1, Seh1) and the TORC1 subunit Kog1, each of which resides on the vacuole membrane and is linked with autophagy through regulation of TORC1 signaling (Figure 2B). To confirm a spatial relationship between Vps501 and the vacuolar interactors, we tested the ability of Vps501 to colocalize with Sea1, Seh1 and Kog1. Our results indicate that Vps501 colocalizes with each vacuolar interactor at the vacuolar membrane (Figure 2C).

**Figure 2.**
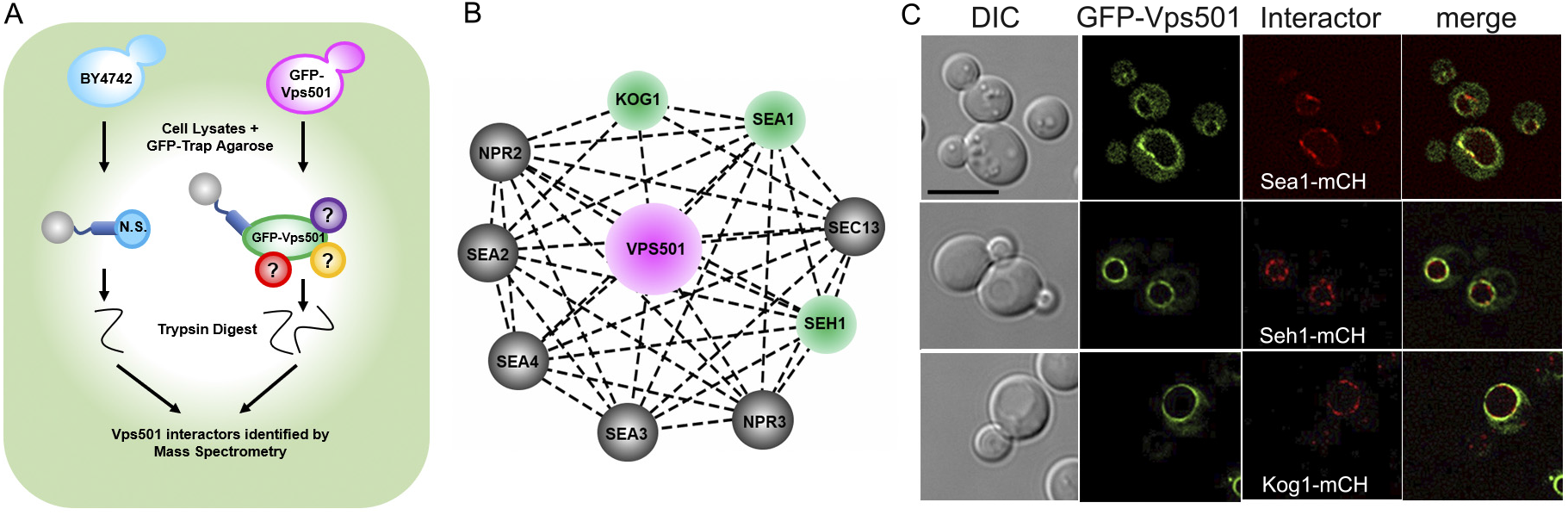
Vps501 interacts with subunits of TORC1 and the SEA complex. (A) Mass spectrometry experimental design to identify Vps501 interactors. GFP-Vps501 and interactors were purified by GFP-Trap affinity purification and SDS-PAGE, followed by in-gel trypsin digestion. Resulting peptides were analyzed and identified by LC–MS/MS according to their relative enrichment. (B) Mass spectrometry analysis using STRING software identified strong interactions with Kog1, a subunit of the TORC1 complex and Sea1 and Seh1, subunits of the SEA complex. Subunits of the SEA complex and TORC1 are represented as nodes in the graphical network. Proteins colored in green (Sea1, Seh1 and Kog1) are those detected by this proteomic study. Lines connecting the nodes represent previously reported interactions. (C) Micrographs show GFP-Vps501 colocalizes with Sea1-mCherry, Kog1-mCherry, Seh1-mCherry at the vacuole membrane. The scale bar indicates 5 μm.

### Vps501-Sea1 interaction stabilizes the SEA Complex and is important for Vps501 vacuolar membrane localization

Due to Vps501’s unique vacuolar localization and apparent interactions with vacuolar proteins, we hypothesized that direct recruitment via the SEA complex is critical for Vps501 vacuolar localization. Indeed, we found that ablation of Sea1 causes mislocalization of GFP-Vps501 to the cytoplasm (Figure 3). Interestingly, deletion of any other SEA complex subunit failed to trigger mislocalization of GFP-Vps501 from the vacuole, suggesting that a interaction with Sea1 specifically mediates Vps501 recruitment to the vacuolar membrane. Among the other Vps501-interacting proteins, we found that the SEACIT subunits Npr2 and Npr3 are dramatically mislocalized to the vacuolar lumen (VL) in 85% of *vps501*Δ*sea1*Δ cells (Figure 4 A-B). While the SEACAT subunits Sea2, Sea3, Sea4 were also mislocalized to the VL in 100% of *vps501*Δ*sea1*Δ cells, this defect was masked by the 80-90% mislocalization observed in the *sea1*Δ single mutant (Supplemental Figure 2). The remaining SEACAT subunits Seh1 and Sec13 were significantly mislocalized from the vacuolar membrane to non-vacuolar compartments in both *sea1*Δ and *vps501*Δ*sea1*Δ cells, but were not found in the vacuole lumen. Seh1 and Sec13 have previously reported roles in the nucleus and ER, respectively, and likely become enriched at these locations when vacuole membrane localization is compromised (Supplemental Figure 2) (Dokudovskaya et al., 2011). Collectively, these results indicate that Vps501 works with Sea1 to stabilize the localization of the SEA complex at the vacuolar membrane, possibly as a part of a larger mechanism that regulates TORC1 signaling.

**Figure 3.**
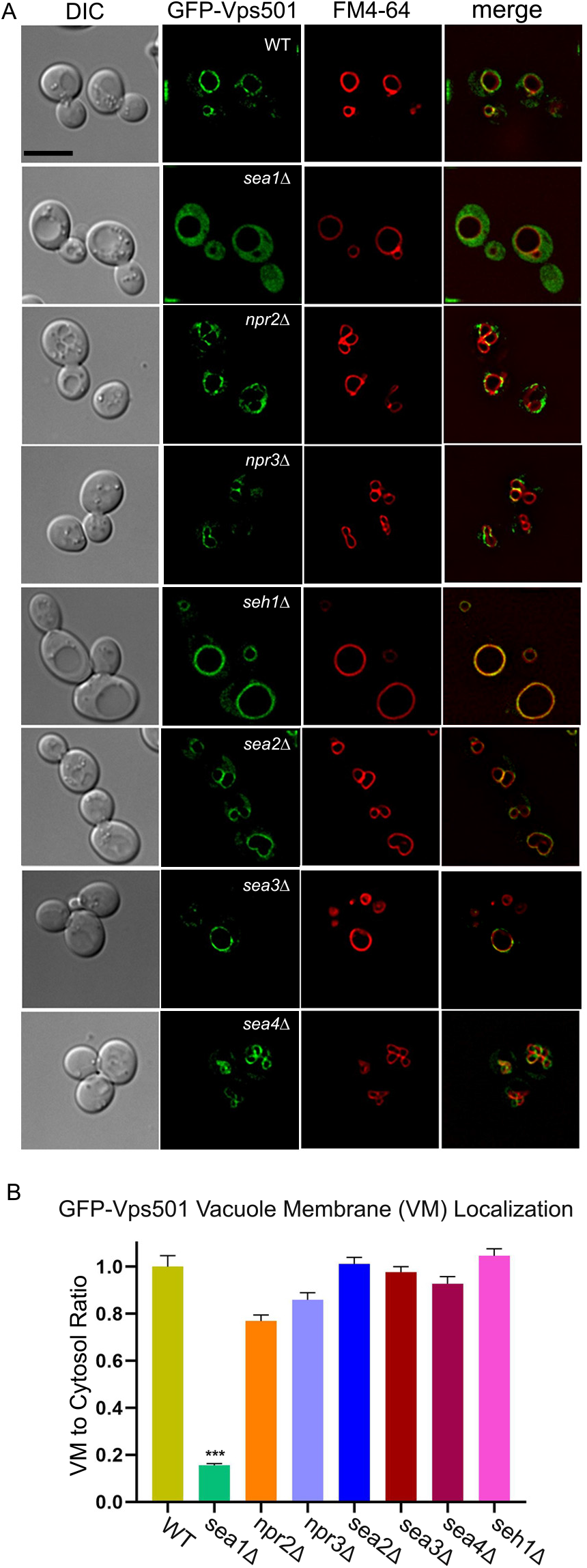
Vps501 requires Sea1 for vacuolar localization. (A) Micrographs of GFP-Vps501 in wildtype and SEA complex mutant cells. GFP-Vps501 is significantly mislocalized in SEACIT mutant cells but not in SEACAT mutant cells. Note, *sec13*Δ cells are nonviable. (B) Vacuole membrane localization was quantified by comparing GFP-Vps501 VML/cytosol ratios in wildtype versus SEA complex mutants. A minimum of 100 cells were measured in triplicate; standard error of the mean was calculated. *p<0.05, ** p<0.01, ***p<0.001 indicates significance as calculated by Student’s t-test. See also supplemental figure 2.

**Figure 4.**
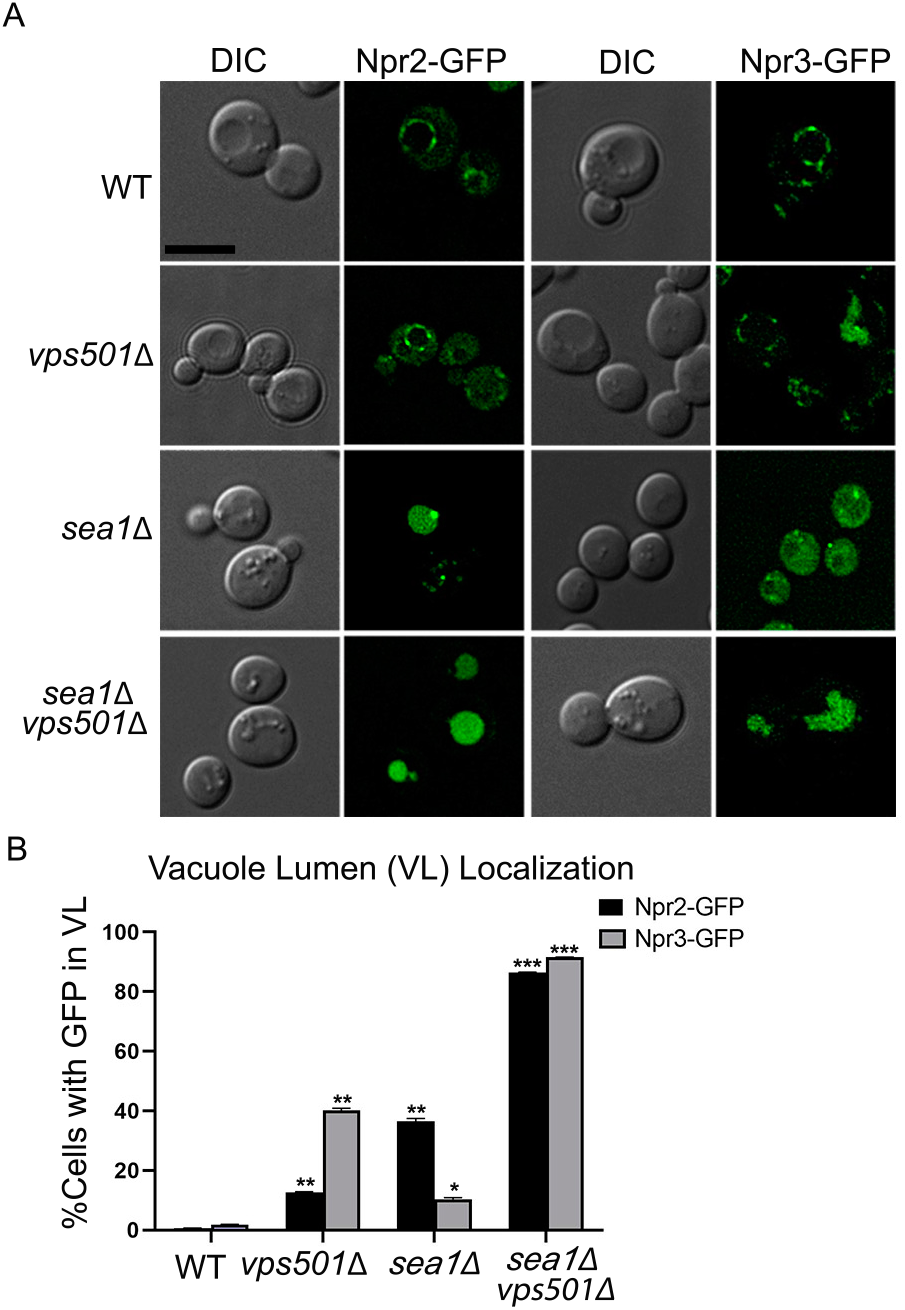
SEACIT subunits, Npr2 and Npr3 require Vps501 and Sea1 for vacuole localization. (A) Npr2-GFP and Npr3-GFP localize to the vacuole membrane in wildtype cells. In *vps501*Δ or *sea1*Δ cells, Npr2-GFP and Npr3-GFP are mislocalized to the vacuole lumen (VL) in 10-40% of cells. In *vps501*Δ*sea1*Δ cells, Npr2-GFP and Npr3-GFP are mislocalized to the VL in 85% of cells. (B) Graph represents vacuole lumen localization (VL) in wildtype cells or mutants, defined by visually scoring the presence of GFP in the vacuole lumen. The results are from three experiments and averaged using the standard error of the mean. Indicated significance is a comparison of wildtype to single deletions or double mutants. *p<0.05, ** p<0.01, ***p<0.001 indicates significance as calculated by Student’s t-test.

### Vps501 contributes to the regulation of autophagy by the SEA complex

We next performed a series of genetic and functional studies to interrogate the potential link between the *VPS501* gene and TORC1 signaling and to determine whether it is epistatic to any of the genes encoding the subunits of the SEA complex. TORC1 signaling can trigger autophagy induction, and we initially used fluorescence microscopy to assess localization of the canonical autophagy marker GFP-Atg8 in strains deleted for *VPS501* singly or in combination with different SEA complex subunits (Figure 5A). GFP-Atg8 processing was also monitored by immunoblotting to detect increases in both total GFP-Atg8 signal and ratio of free GFP to total GFP signal over time (Figure 5B, C). Collectively, these assays allowed us to evaluate whether deletion of *VPS501* results in autophagic flux defects or exacerbates those caused by SEA complex mutants. We found that cells lacking Vps501 display no defects in starvation-induced (nonselective) autophagy, unless combined with deletions of SEACIT subunits. Interestingly, both fluorescence microscopy and immunoblot analysis of *sea1*Δ*vps501*Δ cells showed a striking loss of autophagic flux in GFP-Atg8, indicating that Vps501 and the SEA complex work synergistically during autophagy (Figure 5). Others groups have reported kinetic delays in autophagic flux following deletion of *SEA1*, either alone or in combination with deletion of other SEA complex subunits (Algret et al., 2014). We found that combined loss of *VPS501* and *SEA1* severely impairs autophagy, nearly approximating the complete loss observed in the *atg1Δ* control. It is also worth noting that, as reported in previous studies, *NPR2* and *NPR3* deletions cause significant defects in both bulk and specific forms of autophagy, likely masking any synthetic defect present in *vps501*Δ mutant cells (Dokudovskaya and Rout, 2011). *VPS501* deletion in combination with each of the SEACAT subunits results in a partial reduction in autophagic flux as evidenced by GFP-Atg8 processing defects, with the most significant defect occurring in *seh1Δvps501Δ* cells (Supplemental Figure 3). Given the strong autophagic defects associated with the combined loss of *VPS501* and SEA complex subunits, particularly those of the SEACIT subcomplex, we set out to further examine the role of Vps501 vacuolar membrane localization in autophagy function.

**Figure 5.**
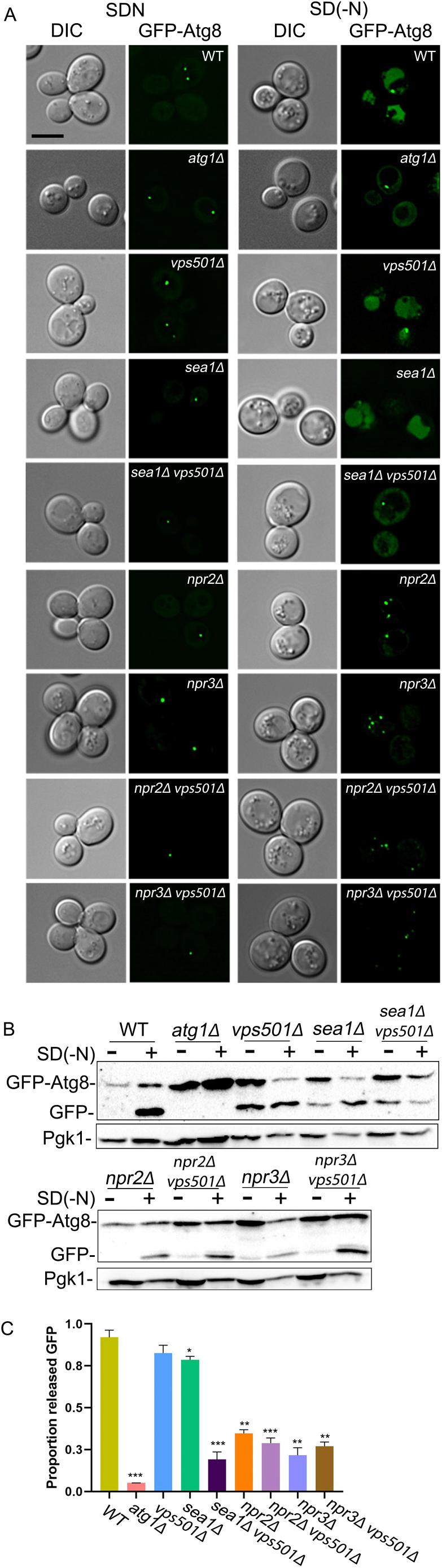
*vps501*Δ*sea1*Δ cells display a synthetic autophagy defect. (A) Maximum projection micrographs of cells expressing GFP-Atg8 in wildtype and indicated mutant cells before and after nitrogen starvation. GFP-Atg8 is found in the vacuole lumen (VL) in wildtype cells after nitrogen starvation, indicating successful autophagic flux has occurred. In *atg1*Δ cells, autophagy induction is inhibited and GFP-Atg8 is absent in the VL. A similar phenotype is also seen only when cells are ablated for both Vps501 and Sea1, indicating a synthetic genetic interaction between Vps501 and Sea1 is critical for autophagy. Note, cells ablated for Npr2 and Npr3 also show major autophagy defects as single knockouts and likely mask any synthetic phenotypes combined with Vps501 ablation. The scale bar indicates 5μm. (B) Quantitative immunoblotting was used to detect the amount of GFP-Atg8 flux before and after autophagy induction. There is a 3-fold decrease in GFP-Atg8 flux in *vps501*Δ*sea1*Δ cells, compared to single deletions, whereas there was no significant difference detected in Npr2 or Npr3 knockout cells. A representative immunoblot is shown. Anti-Pgk1 was used as a loading control. (C) Graph of quantification of GFP-Atg8 processing. The results are from three experiments and averaged using the standard error of the mean. Indicated significance is a comparison of wildtype to single deletions or double mutants. *p<0.05, ** p<0.01, ***p<0.001 indicates significance as calculated by Student’s t-test. See also supplemental figure 3.

### Vps501 contains a non-canonical PX domain that is required for its vacuolar membrane localization and function in autophagy

We next aimed to determine whether the vacuolar membrane localization of Vps501 is required for its autophagic function. In order to generate a mutant Vps501 protein that is unable to localize to the vacuolar membrane, we sought to first identify the domain responsible for this localization. We began by examining the lipid-binding PX domain that might mediate this localization through recognition of the specific lipid composition. Although the canonical lipid-binding motif in the PX domain of Vps501 is missing key motif residues that would be required for PI3P lipid binding, we discovered a predicted secondary noncanonical binding motif (Figure 6A). We used site-directed mutagenesis of the pGFP-Vps501 plasmid to disrupt the noncanonical lipid binding motif in order to determine its effect on localization of Vps501 to the vacuole. We substituted alanine for the key conserved tyrosine and lysine residues in the noncanonical lipid binding motif of Vps501, generating a mutant protein referred to here as GFP-Vps501^YKAA^. Colocalization studies of pGFP-Vps501^YKAA^ and the vacuolar membrane marker FM4-64, indicated that GFP-Vps501^YKAA^ recruitment to the vacuolar membrane is reduced by 70% relative to the GFP-Vps501 control (Figure 6B, D upper graph). This 70% reduction in the vacuolar membrane recruitment of pGFP-Vps501^YKAA^ was seen in both *vps501Δ* and *sea1Δ* cells, and combination of pGFP-Vps501^YKAA^ with *vps501Δsea1Δ* led to a near total loss of Vps501 vacuolar localization (Figure 6B, D upper graph). This indicated that both protein-protein interaction with Sea1 and lipid recognition by the lipid-binding motif are important for Vps501 localization to the vacuolar membrane (Figure 6B, D upper graph). We next ectopically expressed *VPS501YKAA* from a 2-micron plasmid and found that the mutant fails to complement *vps501Δsea1Δ* cells in GFP-Atg8 processing assays (Figure 6C). Taken together with the severe defect in *sea1Δ* cell autophagic flux associated with the *VPS501YKAA* mutant relative to the wild-type control (Figure 6D, upper graph), the combination of Sea1 interaction and lipid-binding appears to be required not only for Vps501 recruitment to the vacuolar membrane but also for its role in autophagy.

**Figure 6.**
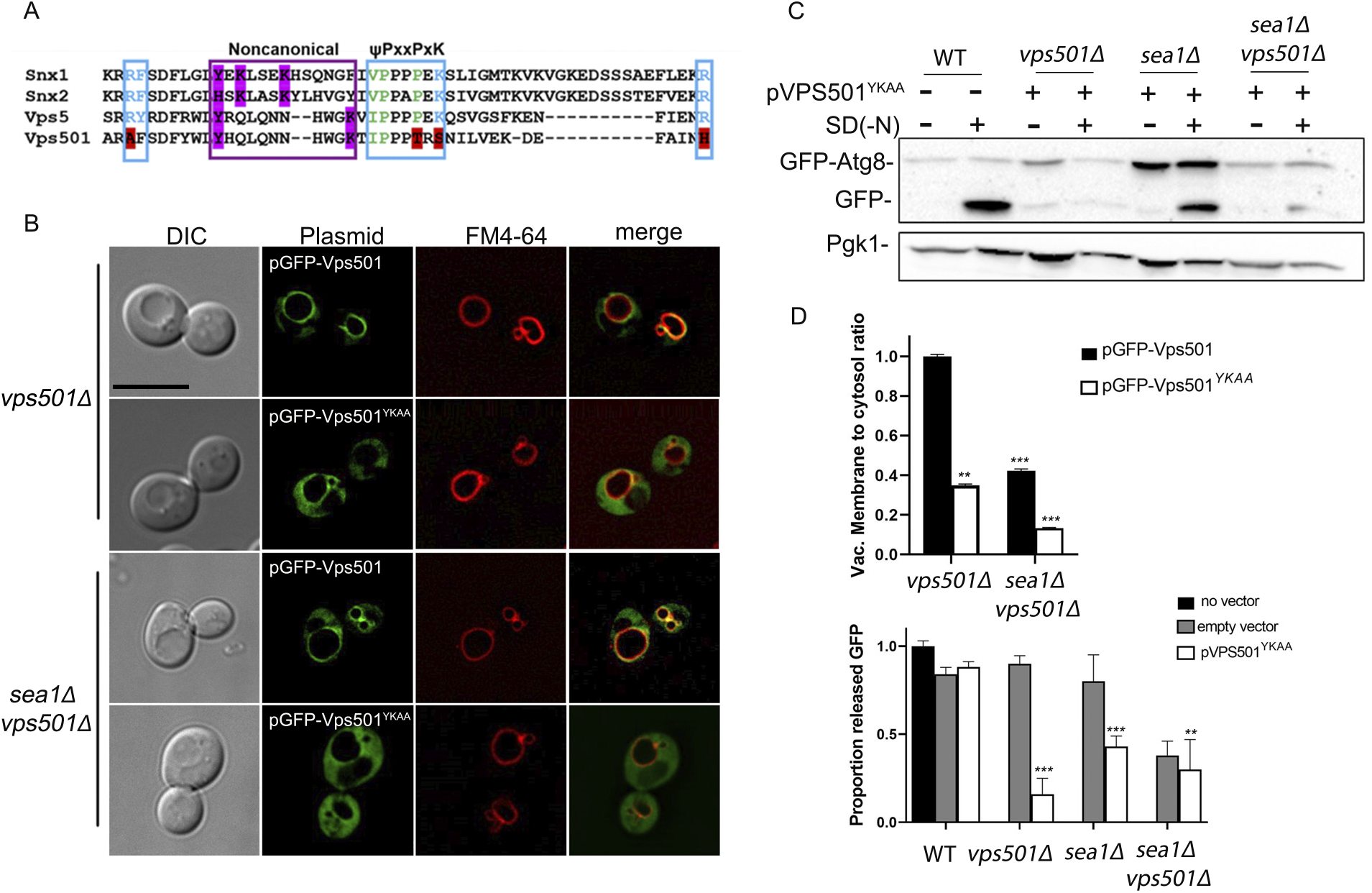
Vps501 non-canonical PX domain is required for localization and function at the vacuole membrane. (A) Sequence alignments of human SNX1, SNX2 and yeast homologs VPS5 and VPS501 PX domains are shown. Blue indicates residues in the canonical PI3P motif. Magenta indicates key secondary lipid binding site residues. Vps501 canonical PI3P motif is not conserved, however, the non-canonical His/Tyr motif is conserved. (B) Alanine mutations to His/Tyr residues in the non-canonical motif resulted in GFP-Vps501^YKAA^ mislocalized to the cytosol in *vps501Δ*, and *vps501Δsea1Δ* cells, indicating Vps501’s recruitment to the vacuole membrane is dependent on both PI3P recognition and Sea1 binding. The scale bar indicates 5μm. (C) Ectopic expression of Vps501^YKAA^ failed to complement *vps501Δsea1Δ* cells in GFP-Atg8 processing assays and severely impairs autophagy flux when expressed in *sea1Δ* cells. (D) Top Graph, quantification of Vps501 localization was defined by vacuole membrane to cytosol ratios as described in material and methods. GFP-Vps501^YKAA^ recruitment to the vacuole membrane is reduced by ~60% in *vps501Δ* cells. When combined with ablation of Sea1 vacuole membrane localization is reduced by ~90%. Bottom Graph, quantification of GFP-Atg8 processing when Vps501^YKAA^ is ectopically expressed in wildtype, *vps501Δ*, or *vps501Δsea1Δ* cells. The results are from three experiments and averaged (standard deviation). Indicated significance is a comparison of wildtype to single deletions or double mutants. *p<0.05, ** p<0.01, ***p<0.001 indicates significance as calculated by Student’s t-test.

### Vps501 and Sea1 regulate Atg27 trafficking to the vacuolar membrane

To further assess the functions of Vps501 in autophagy and regulation of endosomal cargo trafficking, we examined the effects of Vps501 and SEA complex mutations, singly and in combination, on Atg27 localization to the vacuolar membrane. Atg27 is an autophagy-related transmembrane protein and was chosen for investigation as a potential mechanistic link because of its characterized vacuolar-endosomal trafficking itinerary (Segarra et al., 2015). In wild-type cells, Atg27-2XGFP cycles between the vacuolar membrane, Golgi/endosome and autophagic compartments. In *vps501Δsea1Δ* cells, we found that Atg27-2XGFP is significantly depleted from the vacuolar membrane, indicating that Vps501-Sea1 is responsible for maintaining Atg27 at the vacuolar membrane (Figure 7A-B). However, it is not yet clear whether this mechanism is mediated by a direct or indirect interaction with Vps501 and the SEA complex.

**Figure 7.**
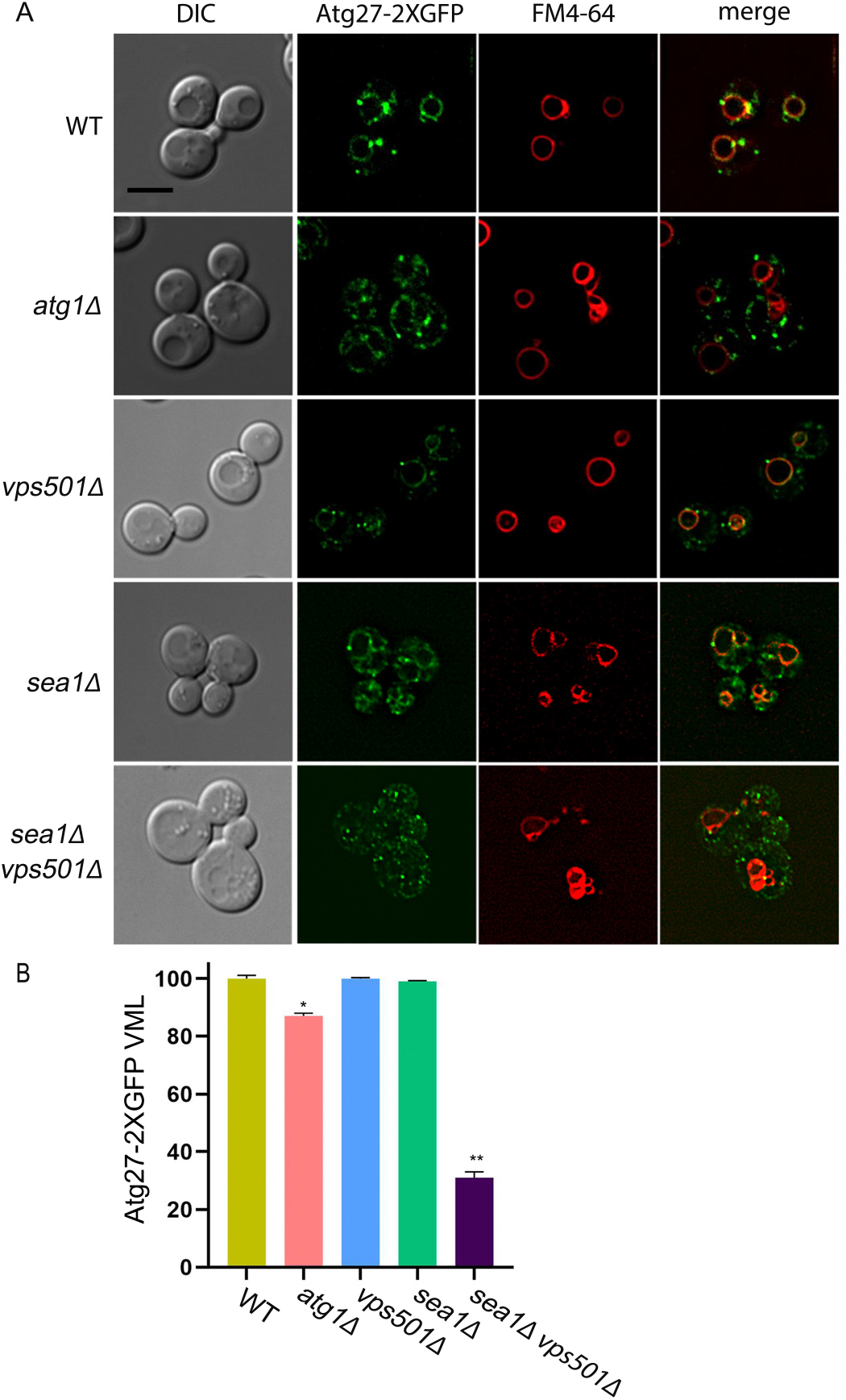
*vps501*Δ*sea1*Δ cells are defective in Atg27 cycling. (A) Micrographs of Atg27-2XGFP in wildtype and indicated mutant cells. Atg27-2XGFP is typically localized to the vacuole membrane, the Golgi and endosomal compartments. In *atg1*Δ cells, Atg27-2XGFP cycling is less abundant on the vacuole membrane. In *vps501Δsea1Δ* cells, Atg27-2XGFP is significantly depleted from the vacuole membrane, indicating Atg27 cycling to and from the vacuole membrane is dependent on Vps501 and Sea1 function during autophagy induction. (B) Graph of the quantification of Atg27-2XGFP vacuolar localization as described in the text. Percentage of cells with Atg27 vacuole localization is between 85-95% in wildtype cells or in cells ablated for only Vps501 or Sea1 and is reduced ~20% *atg1*Δ cells and ~75% in *vps501*Δ*sea1*Δ cells. The results are from three experiments and averaged using the standard error of the mean. Indicated significance is a comparison of wildtype to single deletions or double mutants. *p<0.05, ** p<0.01, ***p<0.001 indicates significance as calculated by Student’s t-test. The scale bar indicates 5μm.

### The TORC1 complex and induction of autophagy are defective in vps501Δsea1Δ cells

While other sorting nexins have previously been found to contribute to autophagy indirectly by mobilizing lipid membranes for autophagosome biogenesis or potentiating vacuolar fusion, the role of Vps501 in autophagy appears to be more direct (Dong et al., 2021; Hanley et al., 2021; Henkel et al., 2020; Ma et al., 2018). The clear physical and genetic interactions of Vps501 with the SEA complex indicate a potential role in regulating TORC1 signaling, possibly during the induction of autophagy. To test this hypothesis, we targeted subunits of TORC1 to determine whether they were defective in v*ps501*Δ*sea1*Δ cells. We were particularly interested in the Kog1 subunit, as it was one of our most abudant hits in the proteomics screen for Vps501 interactors (Figure 2). Similar to what others have found, we localized Kog1-2XGFP to two independent TORC1 pools; one around the vacuolar membrane and a second in dot-like perivacuolar structures (Chen et al., 2021; Hatakeyama et al., 2019)(Figure 8A). This dual localization is consistently present in wildtype, *vps501*Δ *or sea1*Δ cells. Interestingly, the dot-like perivacuolar structures exhibit enhanced accumulation in *vps501*Δ*sea1*Δ cells, with the average number of Kog1-2XGFP dots enriched 2-fold relative to the wildtype or single mutant controls. The vacuolar Kog1-2XGFP pool appears be reduced, either in the presence or absence of nitrogen (Figure 8A-B). Recently, these dot-like TORC1 structures have been referred to as signaling endosomes and have been shown to have unique phosphorylation targets (Chen et al., 2021; Hatakeyama et al., 2019). One such target is Atg13, a regulatory subunit of the Atg1 signaling complex that is required for induction of autophagy (Kamada et al., 2000). When TORC1 is active, Atg13 is phosphorylated, inhibiting induction of autophagy. Therefore, we hypothesized that Atg13 phosphorylation would be defective in *vps501*Δ*sea1*Δ cells if autophagy induction were impaired. We analyzed Atg13 by immunoblotting in wildtype and mutant cells, both before and after nitrogen starvation, and quantified the resulting signals. In wildtype, *vps501*Δ *or sea1*Δ cells, the majority of Atg13 is phosphorylated as indicated by an enrichment of the upper Atg13 band (labeled Atg13P) during vegetative growth, indicating that TORC1 is active (Figure 8C-D). Autophagy-inducing conditions inactivate TORC1, and the enrichment of the lower band (labeled Atg13) in wildtype, *vps501*Δ, *or sea1*Δ cells indicates that the majority of Atg13 is not phosphorylated (Figure 8C-D). In contrast, Atg13 appears only phosphorylated in *vps501*Δ *sea1*Δ cells, regardless of nitrogen starvation, indicating that autophagy induction is defective in these mutants, (Figure 8C-D). Taken together, these results indicate that Vps501 cooperates with the SEA complex to regulate TORC1 signaling during the induction of autophagy.

**Figure 8.**
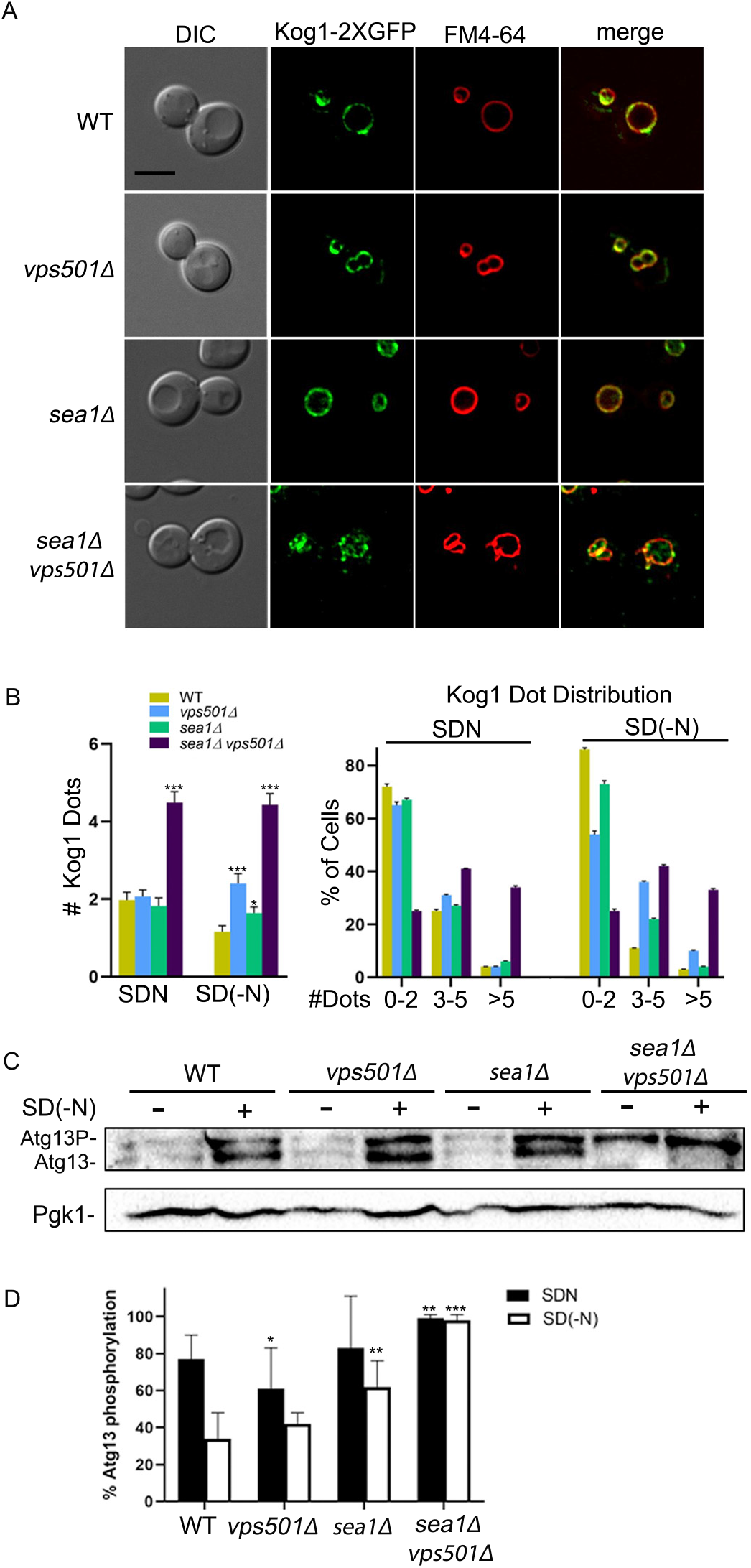
TORC1 and autophagy induction is defective in *vps501*Δ*sea1*Δ cells. (A) Micrographs of Kog1-2XGFP localization in wildtype and indicated mutant cells during vegetative growth. Kog1 a subunit of TORC1, typically localizes to the vacuole membrane and in dot-like structures juxtaposed the vacuole membrane, known as signaling endosomes. In *vps501*Δ*sea1*Δ cells, these dot-like structures accumulate indicating the pool of signaling endosomes are enriched, while the vacuole TORC1 pool appears reduced. Vacuolar membranes are shown using FM4-64 dye. (B) The number of Kog1 dot-like structures were quantified in wildtype and indicated mutant cells before and after nitrogen starvation as described in the text. Regardless of starvation conditions, there was a 2-fold increase in dot-like structures in *vps501*Δ*sea1*Δ cells as compared to single mutants or wildtype cells (left graph). 3 or more dot-like structures were found in 80% of *vps501*Δ*sea1*Δ cells, while single mutants or wildtype cells typically had 0-2 dots (right graph). *p<0.05, ** p<0.01, ***p<0.001 indicates significance as calculated by a one-way ANOVA from three biological replicates. (C) Quantitative imunnoblot analysis of Atg13 in wildtype and indicated mutant cells before and after nitrogen starvation. In wildtype, *vps501*Δ *or sea1*Δ cells, the majority of Atg13 is phosphorylated as indicated by an enrichment of the upper Atg13 band (labeled Atg13P) during vegetative growth, indicating TORC1 is active. In autophagy induction conditions, TORC1 is inactivated and the majority of Atg13 is not phosphorylated indicated by an enrichment of the lower band (labeled Atg13). In *vps501*Δ *sea1*Δ cells, Atg13 appears only phosphorylated indicating autophagy induction is defective in these mutants, regardless of nitrogen starvation. (D) Percentage of Atg13 phosphorylation was quantified by determining the proportion of Atg13P to total protein using densitometry. A representative immunoblot is shown. Anti-Pgk1 was used as a loading control. Indicated significance is a comparison of wildtype to single deletions or double mutants. *p<0.05, ** p<0.01, ***p<0.001 indicates significance as calculated by Student’s t-test from three biological replicates.

## Discussion

In this study, we report the identification of a novel vacuolar membrane SNX-BAR protein, Vps501. While phylogenetically, Vps501 is most related to Vps5, an essential component of the yeast retromer complex, protein sequence analysis shows residue variations acquired during its divergence (Figure 1, Supplemental Figure 1). In this study, we demonstrate how divergence from Vps501 and Vps5 has resulted in unique differences in function. For example, current models indicate that an evolutionarily conserved Phox-homology (PX) domain, a feature common in all sorting nexins, binds specifically to PI3P, the major phospholipid of the endosome. However, Vps501 has no canonical PI3P binding motif, as it is missing key motif residues. Instead, Vps501 has a secondary site that is likely solely responsible for lipid binding. Also, Vps501 exclusively resides on the vacuolar membrane, a unique feature amongst the SNX-BAR protein family. We speculate Vps501 likely possesses lipid specificity beyond PI3P as was discovered in other sorting nexins (Chandra et al., 2019). This notion was supported by the Vps501^YKAA^ mutant mislocalizing to the cytosol, leaving only a small percentage remaining on the vacuolar membrane (Figure 6). Interestingly, in *sea1Δ vps501YKAA* cells, the cytoplasmic mislocalization of the Vps501^YKAA^ mutant was completely blocked from the vacuolar membrane, suggesting Vps501 localization requires both the vacuolar membrane lipid composition and resident vacuolar proteins. Whether these two binding requirements are universal or specialized to each SNX-BAR protein is not known. However other SNX-BAR proteins such as Mvp1, an endosomal SNX-BAR, was recently found to tetramerize and dissociate in order to bind membrane, indicating a selective combination of protein-protein interaction regulation and lipid specificity likely determine the function of each SNX-BAR (Sun et al., 2020). Understanding the novel ways in which Vps501 appears to recognize vacuolar lipid will be critical to understanding its function and regulation.

Multiple lines of evidence support a role for Vps501 in TORC1 signaling during autophagy induction. Firstly, using a co-immunoprecipitation mass spectrometry approach, we identified subunits of the evolutionary conserved SEA complex (Sea1, Seh1) and TORC1 subunit Kog1 (Figure 2–3). Each of these identified proteins resides on the vacuolar membrane and colocalizes with Vps501. Furthermore, Sea1 was found to be critical for Vps501 localization, while the SEA complex was destabilized in *vps501*Δ *sea1*Δ cells, indicating a direct interaction (Figure 3, Supplemental Figure 2). It is also worth noting that the previous studies in which mass spectrometry-based proteomics were used to discover the eight components of the SEA complex all utilized an individual band excision-based proteomic approach focusing on the most enriched bands (Dokudovskaya and Rout, 2011). This design excluded low abundant or low affinity interactions and may account for why Vps501 was not detected in their screens.

Secondly, we found that Vps501 works together with the SEA complex to mediate GFP-Atg8 autophagic flux. We found that cells lacking Vps501 display a severe deficiency in autophagy only when SEA complex subunits were deleted as well (Figure 5, Supplemental Figure 3). A recent study came to similar conclusions, showing that loss of Sea2, Sea3 or Sea4 did not trigger major defects unless combined with *sea1Δ* (Algret et al., 2014; Dokudovskaya and Rout, 2011; Dokudovskaya et al., 2011). This led us to believe that Vps501 and the SEA complex cooperate within a synergistic pathway during autophagy induction. Furthermore, the complete impairment of autophagic flux in *vps501YKAA* cells suggests that a negative mutation within the vacuolar membrane recognition site of Vps501 is sufficient to drive impairment of the SEA complex.

Lastly, we have defined the role of Vps501 as a co-regulator of autophagy that promotes SEACIT inhibition of TORC1 during autophagy induction. The yeast Rag GTPase-TORC1 complex is found in two spatially and functionally distinct pools, on both the vacuolar and endosomal membranes (Chen et al., 2021; Hatakeyama et al., 2019). Vacuolar TORC1 promotes protein synthesis through its proximal effector Sch9, while endosomal TORC1 controls autophagy induction through phosphorylation of Atg13, preventing Atg1 complex formation at the pre-autophagosomal structure. Interestingly, *vps501*Δ*sea1*Δ cells show depletion of the Kog1-2XGFP subunit of TORC1 from the vacuolar membrane and enrichment in dot-like structures by nearly 2-fold (Figure 8). Likewise, the endosome-specific TORC1 substrate Atg13 is hyperphosphorylated in *vps501*Δ *sea1*Δ cells (Figure 8). This confirms that Vps501 function is specific to the vacuolar pool of TORC1 and indicates that failure to induce autophagy is the underlying defect in *vps501*Δ*sea1*Δ cells. In addition, the autophagy defects observed in *vps501*Δ *sea1*Δ cells were specifically observed upon autophagy induction through nitrogen deprivation and not through rapamycin treatment. Briefly, we found that *vps501*Δ *sea1*Δ mutant cells did not display autophagy influx defects when they received rapamycin treatment. Rapamycin is a potent lipophilic macrolide antifungal drug that inhibits TORC1 signaling and is widely used to induce autophagy in yeast (Barbet et al., 1996; Rohde et al., 2001). This suggests that Vps501 and Sea1 are likely operating upstream of TORC1, allowing rapamycin to bypass the autophagic defects of the *vps501*Δ *sea1*Δ mutant.

We also found that the Vps501-SEA complex mediates Atg27 trafficking from the vacuolar membrane. We do not believe that autophagy triggers this phenomenon directly, since autophagy-deficient *atg1Δ* cells display only a minor loss of Atg27 trafficking. Instead, we believe that Vps501-SEA control over Atg27 trafficking may be an indirect consequence of TORC1 hyperactivation at the signaling endosome. Further studies are needed to test this hypothesis.

Collectively, these findings lead us to propose a new model for Vps501 function at the vacuolar membrane (Figure 9). When wildtype cells are deprived of nitrogen, the SEA complex acts through effector molecules such as the EGO complex to inhibit TORC1 and facilitate autophagic induction. Vps501 acts as a structural stabilizer to tether the SEA complex to the vacuolar membrane, relying on its direct interaction with Sea1 and the lipid interactions of its non-canonical PX domain to facilitate the induction of autophagy and proper Atg27 trafficking. In contrast, *vps501*Δ *sea1*Δ cells exhibit destabilization of the SEA complex, causing TORC1 to remain active under nitrogen-starved conditions. This results in defects both in the induction of autophagy and in Atg27 trafficking.

**Figure 9.**
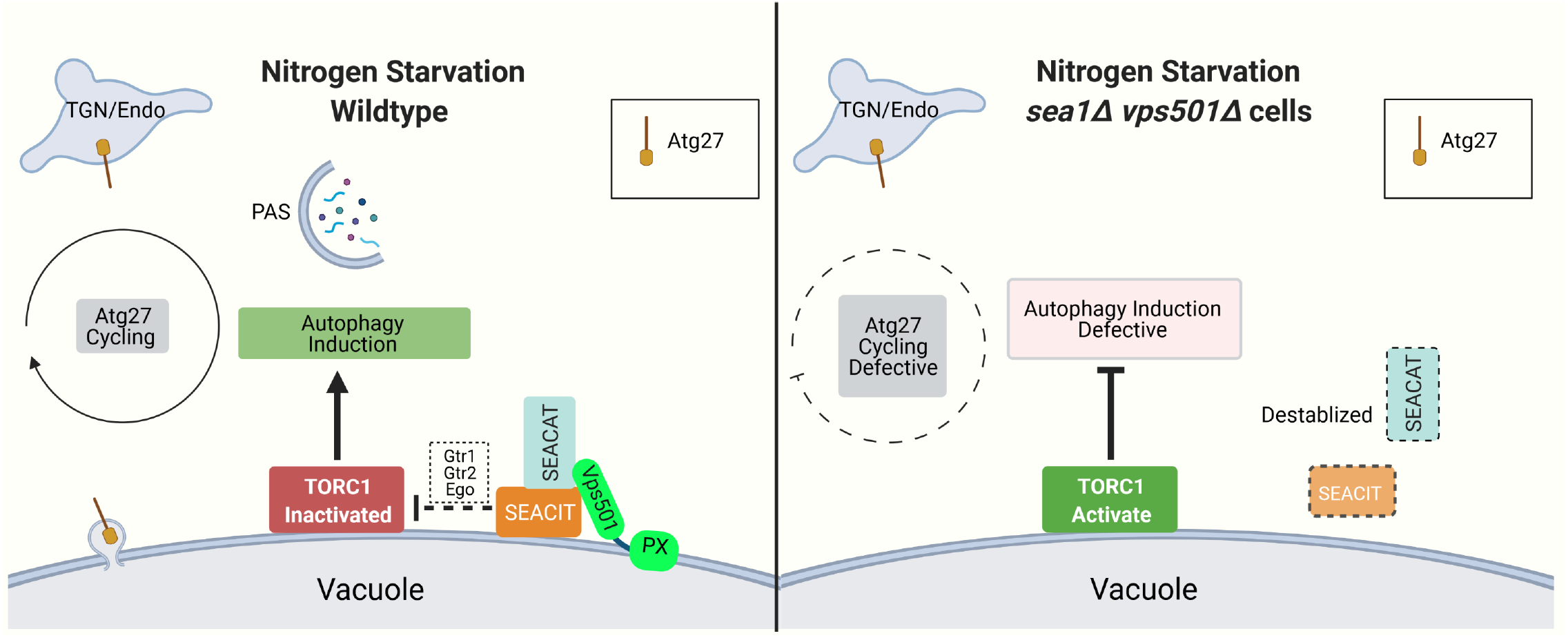
Vps501 cooperates with the SEA complex to inactivate TORC1. Under starvation conditions, Vps501 stabilizes the SEA complex to the vacuole membrane using a non-canonical PX domain and a direct interaction with Sea1. This interaction promotes SEACIT inactivation of TORC1 via the EGO/GTR complex, resulting in the induction autophagy with proper Atg27 cycling. In *vps501*Δ *sea1*Δ cells, the SEA complex is destabilized and TORC1 remains activate, resulting in defective autophagy induction and defective Atg27 cycling.

## Materials and Methods

### Phylogenetic Analysis

To determine the identity of Saccharomyces cerevisiae YKR078W in relation to other SNX-BAR sorting nexins, we conducted a phylogenetic analysis. We used a former analysis of SNX-BAR proteins in opisthokonts (e.g., fungi Vps5, and Vps17, animal and choanoflagellate SNX 5/6/32 and SNX 1/2) by Koumandou et al. (2011) as an initial seed alignment. We added additional Vps5 proteins from other fungal species from the family Saccharomycetaceae for better resolution of fungal Vps5 proteins, including YKR078W. Furthermore, in order to identify if other fungi have a protein similar to YKR078W, we used BLASTp to query protein models at FungalDB to identify additional SNX-BAR sorting nexins with similarity to YKR078W from S. cerevisiae. These searches only recovered the single Vps5 protein from the fungal species. No additional proteins were detected with an E-value less than 10 (search threshold). Full-length sequences for all taxa for each gene family were aligned with Muscle 3.6 (Edgar, 2004) and edited manually in the case of clear errors. Maximum likelihood analyses were conducted with RAxML v.8.2.4 (Stamatakis, 2014) using a LG+G matrix model determined by ProtTest v.3, (Darriba et al., 2011) and a trimmed alignment containing the conserved PX +BAR and SNX-BAR domains. Support for particular nodes for maximum likelihood analyses was assessed with 1,000 bootstraps. Trees were visualized and illustrated with FigTree v1.4 (http://tree.bio.ed.ac.uk/software/figtree/).

### Yeast Strains and Culture Conditions

Yeast strains were grown using standard media and conditions (Guthrie and Fink, 1991) unless indicated. Yeast strains were constructed in BY4742 (*MAT*α *his3-1*, *leu2-0*, *met15-0*, and *ura3-0)* by homologous recombination of gene-targeted, polymerase chain reaction (PCR)-generated DNAs using the method of Longtine et al. (Longtine et al., 1998) and/or derived from the EUROSCARF KANMX deletion collection (Open Biosystems/Thermo Scientific, Waltham, MA) or produced by replacement of the complete reading frame with the *HIS3MX6* or *URA3* cassette. Gene deletions were confirmed by PCR amplification of the deleted locus. Cells were grown in standard synthetic complete medium lacking nutrients required to maintain selection for auxotrophic markers and/or plasmid, unless indicated otherwise. To induce bulk or non-selective autophagy, cells were grown to log phase, harvested, and transferred to SD(-N) medium for nitrogen starvation (2% dextrose, 0.17% Yeast Nitrogen Base without amino acids and without ammonium sulfate) for 16 hours.

For the construction of an integrated N terminal GFP-Vps501 yeast strain under a GPD promoter, we PCR amplified pGFP-^GPD^Vps501 (described below) using the following primers that included 50 bp flanking the *VPS501*locus; ATCAGAACTGCAACCCTACAGATTAGATATGGAGAACGACAAGGCGTCACGTGAGCAAGGGCGAGGAGCTGTTCA and GCTTTTTCAGTAGTAAATTATCTTCTTTAATTACGTTATTATGTACATATTTGGCTTATGTGCTCATCTGGTACA). PCR products were subsequently transformed into cells ablated for *VPS501* (by replacement with a URA marker) and allowed to homologously recombine into the *VPS501* locus. Resulting clones were selected on F.O.A 5-Fluoroorotic Acid (5-FOA) and confirmed by PCR.

### Light Microscopy and Image Analysis

Yeast cells from cultures grown to OD_600_ ≈ 0.5 were mounted in growth medium, and 3D image stacks were collected at 0.3-μm z increments on a DeltaVision elite workstation (Cytiva) based on an inverted microscope (IX-70; Olympus) using a 100×1.4NA oil immersion lens. Images were captured at 24°C with a 12-bit charge-coupled device camera (CoolSnap HQ; Photometrics) and deconvolved using the iterative-constrained algorithm and the measured point spread function. To visualize vacuole morphology, yeast cells were labeled with 7-Aminochloromethylcoumarin (CMAC; Life Technologies) at a concentration of 100 μM for 30 min in synthetic medium at room temperature. To visualize the vacuole membrane, FM4-64 (32nM) was added to cell cultures for 20 min at 30°C. Cells were then washed, resuspended in fresh medium, and then incubated for 60 minutes to allow FM4-64 to accumulate in the vacuole membrane adapted from (Vida and Emr, 1995). Image analysis and preparation was done using Softworx 6.5 (Cytiva) and ImageJ v1.50d (Rasband).

The analysis of vacuole membrane localization for GFP-Vps501, GFP-Vps501^YKAA^ and Atg27-2XGFP was determined using a manual method implemented using ImageJ v1.53c (Rasband). A region of interest (ROI) was selected to contain a single cell and the total sum of GFP fluorescence was calculated (TF). Next, we used the Mask macro to delineate the vacuole ROI defined by FM4-64 and overlayed onto the GFP channel to define the vacuole fluorescence (VF). To calculate cytosol fluorescence intensity (CF), the vacuole mask was inverted so that all pixels outside of the mask were assigned a maximum value and the regions corresponding the vacuole signal were assigned a value of zero. A ratio of the VF to TF is presented in the graphs. Vacuole membrane localization for Seh1-GFP, Sec13-GFP and Atg27-2xGFP, were calculated by calculating the percent of cells with GFP signal on the vacuole using FM4-64 and CMAC as visual guides to determine vacuole boundaries. To quantify vacuole lumen localization, wildtype cells or mutants were visually scored for presence of GFP in the vacuole lumen. Kog1-GFP puncta numbers were quantified from z stacks collected at 0.3-μm intervals. Total patches/cell were counted from maximum intensity projections from small budded cells. A minimum of 100 cells was used in all experimental conditions and performed in biological triplicate.

### GFP-Atg8 Processing and Immunoblotting

For quantitative immunoblot analysis of GFP-Atg8, cells were grown under standard vegetative or autophagy inducing conditions to OD_600_ ≈ 0.5, as described above. Typically, 3.0 x 107 cells were harvested by centrifugation and lysed by glass bead agitation in SDS-PAGE sample buffer. 10% polyacrylamide gels were loaded with 5.0 X107 cell equivalents and transferred onto standard 0.45 μm nitrocellulose. Anti-GFP primary mouse monoclonal antibody (1814460, Roche) was diluted 1:2500 and Santa Cruz (sc-2055) goat anti-mouse HRP-conjugated antibody was used at 1:10000. Anti-Pgk1 at 1:5000 (Life Technologies) was used as loading controls. Centromeric GFP-Atg8 (Shintani et al., 2002) plasmids were used in the processing assays.

Atg13 immunoblots were done as previously described (Kamada, 2017) with the following modifications. 3.0 x 10^7^ cells were harvested by centrifugation and precipitated by Trichloroacetic acid (TCA). 7.5% polyacrylamide gels were loaded with 0.75 X10^7^ cell equivalents. Anti-Atg13 primary rabbit antibody was generously provided by Dr. Yoshiaki Kamada and used at 1:5000 and goat anti-rabbit HRP-conjugated antibody was used at 1:5000. p3xHA-Atg13 was purchased from Addgene (Plasmid #59544) and used in all indicated experiments. All enhanced chemiluminescence *(ECL)* blots were development on a Chemidoc-MP (Bio-Rad) and band intensities were quantified using Quantity One 1D analysis software (Bio-Rad) and all statistical analysis done using GraphPad Prism 8.

### Plasmids

pGFP-^GPD^Vps501 was constructed using Gateway cloning. Plasmid insert was made using PCR amplified wild-type genomic *VPS501* locus with the following primers: GGGGACAAGTTTGTACAAAAAAGCAGGCTTAGAGAACGACAAGGCGTCACAT and GGGGACCACTTTGTACAAGAAAGCTGGGTTTCATTGGCTTATGTGCTCATCTGGT and cloned using a BP recombination reaction into pDONR221. Resulting DONOR vector was recombined with pAG425GPD-EGFP-ccDB (Alberti et al., 2007) in a final LR recombination reaction to generate the pAG415GPD-eGFP-Vps501 expression clone (pGFP-^GPD^Vps501).

pGFP-Vps501^YKAA^ was made commercially using site-directed mutagenesis (GenScript, Piscataway, NJ), introducing alanine mutations into following positions Y160A and K170A of pGFP-^GPD^Vps501. pVps501^YKAA^ was derived from pGFP-Vps501^YKAA^ using Gateway cloning as described above.

### Mass spectrometry

Yeast strains expressing GFP-Vps501 were grown to 0.5 X107 cell density in YPD media. GFP fusion proteins were purified as follows: 200mg of protein extract was incubated with GFP-Trap Magnetic Agarose (ChromoTek) at 4°C for 20 minutes with gentle agitation. GFP-Trap beads were collected and washed five times (50mM Tris-HCL pH 7.4, 150mM NaCl, Roche complete Protease Inhibitor Cocktail EDTA free). After the final wash, the buffer was aspirated and the GFP-Trap beads were incubated with 5x SDS-PAGE sample buffer and denatured for five minutes at 95°C. 20 μL of each sample was analyzed by SDS-PAGE, followed by an immunoblot. The experiment was performed in triplicate and normalized to the Nano-Trap magnetic beads alone.

### Trypsin digestion of samples from SDS-PAGE gel plugs

The gel plugs for each sample were excised by a sterile razor blade, divided into 2 sections 1 cm each, and additionally chopped into 1 mm pieces. Each section was washed in dH2O and destained using 100 mM NH_4_HCO_3_ pH 7.5 in 50% acetonitrile. A reduction step was performed by addition of 100 ml 50mM NH_4_HCO_3_ pH 7.5 and 10 ml of 200mMtris(2-carboxyethyl) phosphine HCl at 37°C for 30 min. The proteins were alkylated by addition of 100 ml of 50 mM iodoacetamide prepared fresh in 50 mM NH_4_HCO_3_ pH 7.5 buffer, and allowed to react in the dark at 20°C for 30 min. Gel sections were washed in water, then acetonitrile, and vacuum dried. Trypsin digestion was carried out overnight at 37°C with 1:50-1:100 enzyme-protein ratio of sequencing grade-modified trypsin (Promega) in 50 mM NH_4_HCO_3_ pH 7.5, and 20mM CaCl2. Peptides were extracted with 5% formic acid and vacuum dried and sent to the Mayo Clinic Proteomics Core facility for HPLC and LC-MS/MS data acquisition.

### HPLC for mass spectrometry

All samples were re-suspended in Burdick & Jackson HPLC-grade water containing 0.2% formic acid (Fluka), 0.1% TFA (Pierce), and 0.002% Zwittergent 3-16 (Calbiochem), a sulfobetaine detergent that contributes the following distinct peaks at the end of chromatograms: MH^+^ at 392, and in-source dimer [2 M+ H^+^] at 783, and some minor impurities of Zwittergent 3-12 seen as MH^+^ at 336. The peptide samples were loaded to a 0.25 ml C8 OptiPak trapping cartridge custom-packed with Michrom Magic (Optimize Technologies) C8, washed, then switched in-line with a 20 cm by 75 mmC18 packed spray tip nano column packed with Michrom Magic C18AQ, for a 2-step gradient. Mobile phase A was water/acetonitrile/formic acid (98/2/0.2) and mobile phase B was acetonitrile/isopropanol/water/formic acid (80/10/10/0.2). Using a flow rate of 350 nL/min, a 90 min, 2-step LC gradient was run from 5% B to 50% B in 60 min, followed by 50%-95% B over the next 10 min, hold 10 min at 95% B, back to starting conditions and re-equilibrated.

### LC-MS/MS analysis

Electrospray tandem mass spectrometry (LC-MS/MS) was performed at the Mayo Clinic Proteomics Core on a Thermo Q-Exactive Orbitrap mass spectrometer, using a 70,000 RP survey scan in profile mode, m/z 340-2000 Da, with lock masses, followed by 20 MSMS HCD fragmentation scans at 17,500 resolution on doubly and triply charged precursors. Single charged ions were excluded, and ions selected for MS/MS were placed on an exclusion list for 60 s.

### LC-MS/MS data analysis, statistical analysis

All LC-MS/MS *.raw Data files were analyzed with MaxQuant version 1.5.2.8, searching against the SPROT Saccharomyces cerevisiae database downloaded 09.28.2017 and searched using the following criteria: LFQ quantification with a min of 1 high confidence peptide. Trypsin was selected as the protease with max miss cleavage set to 2. Carbamidomethyl (C) was selected as a fixed modification. Variable modifications were set to Deamidation (NQ), Oxidization (M), Formylation (n-term), and Phosphorylation (STY). Orbitrap mass spectrometer was selected using a MS error of 20 ppm and a MS/MS error of 0.5 Da. A 1% FDR cutoff was selected for peptide, protein, and site identifications. LFQ Intensities were reported based on the MS level peak areas determined by MaxQuant and reported in the proteinGroups.txt file. Proteins were removed from the results file if they were flagged by MaxQuant as “Contaminants”, “Reverse” or “Only identified by site”. Complete three biological replicates were performed. The abundance data from each biological replicate were normalized to the ratio of Vps501 bait protein in that biological replicate. LFQ Peak intensities were analyzed in each run to determine protein hits that fell into the category of either Vps501 elution (VE) only hits or Bead elution (BE) only hits and retained if they confirmed to VE state across all 3 runs. LFQ Sig cutoffs are Sig Up > 1.2 ratio (Log2 0.26) and Sig Down < 0.8 ratio (Log2 −0.32). Any hits that were not observed in at least 2 replicates each were labeled ‘no quant’ (a normalized ratio was still calculated but not included in final data set analysis). A list of proteins identified and corresponding ratios can be found in Supplemental Materials. The mass spectrometry proteomic data have been deposited to the ProteomeXchange Consortium (http://proteomecentral.proteomexchange.org) (Deutsch et al., 2017) via the PRIDE partner repository (Vizcaino et al., 2013; Vizcaino et al., 2016).

## Acknowledgements

We are grateful to Mandi Ma and members of the Chi, Truman, and Reitzel Labs for thoughtful discussions and comments on this manuscript. This work was supported by the National Science Foundation 2028519 to RJC, National Institutes of Health award 1R01GM139885 to AWT, and UNC Charlotte Faculty Research Grants Program to RJC and AMR.

## Abbreviations List

BAR: Bin-Amphiphysin-Rvs161 (BAR) homology
CBB: coomassie brilliant blue
GFP: Green Fluorescent Protein
ORF: Open reading frame
PGK: phosphoglycerate kinase
PX: Phox homology domain
PI3P: phosphatidylinositol-3-phosphate
ROI: region of interest
SNX: sorting nexin
SNX-BAR: sorting nexin containing a BAR domain
WT: wild-type
TEN: Tubular Endosomal Network
VL: Vacuolar Lumen
VM: Vacuolar Membrane
WT: wild-type

---

**Supplemental Figure 1.**
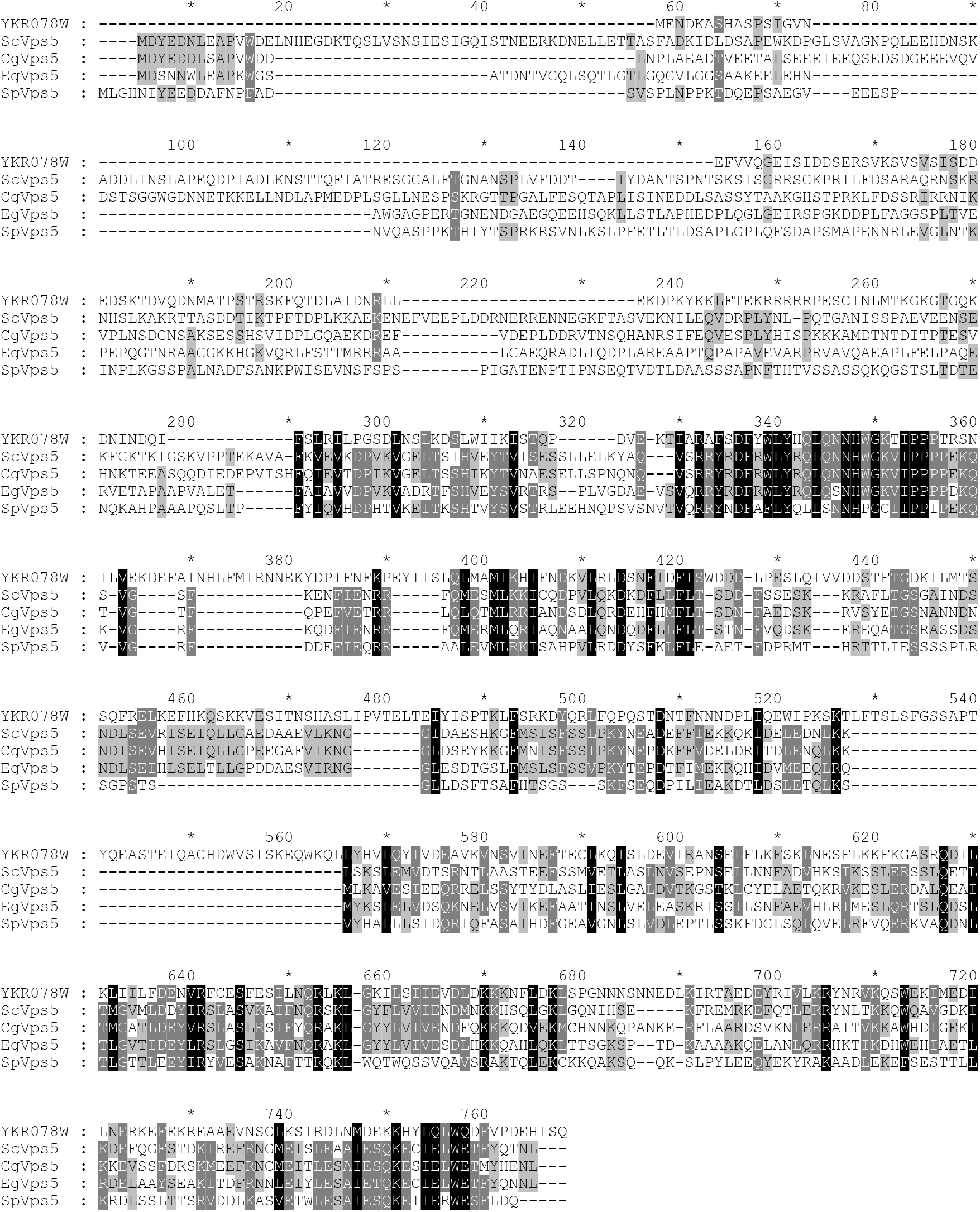
Sequence alignment of YKR078W/VPS501 to Vps5 proteins from other yeast taxa. Full-length sequences for all taxa for each gene family were aligned with Muscle 3.6 (Edgar, 2004) and edited manually in the case of clear errors. Maximum likelihood analyses were conducted with RAxML v.8.2.4 (Stamatakis, 2014) using a LG+G matrix model determined by ProtTest v.3 (Darriba et al., 2011) and a trimmed alignment containing the conserved PX-BAR domains. Sc, *S. cerevisiae*, Cg, *C. glabrata*, Eg, *E. gossypi*, Sp, *S. pombe*.

**Supplemental Figure 2.**
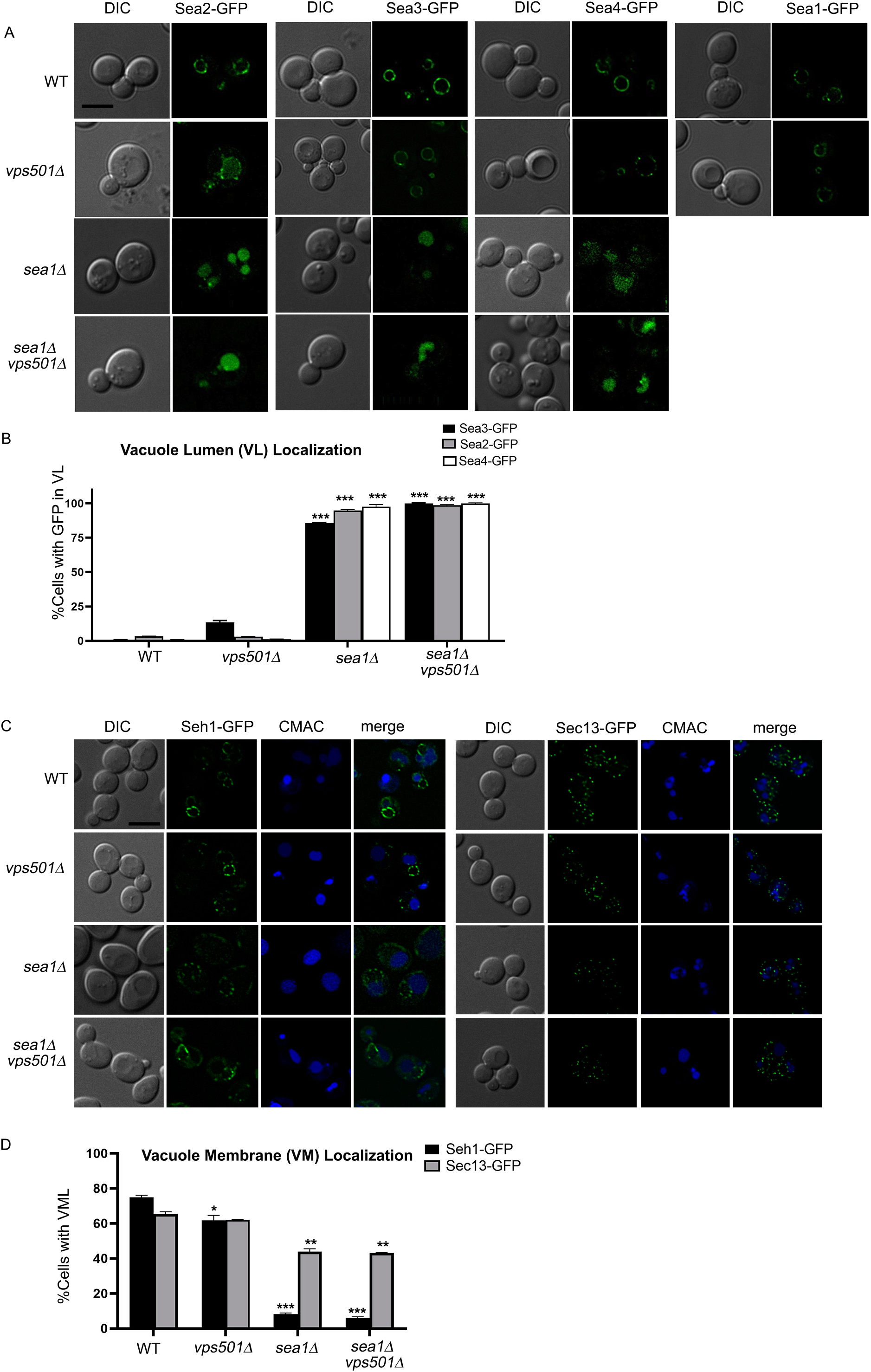
SEACAT subunits, Sea2-GFP, Sea3-GFP, Sea4-GFP, Seh1-GFP and Sec13-GFP are partially mislocalized in *vps501*Δ*sea1*Δ cells. (A) Sea2-GFP, Sea3-GFP, Sea4-GFP and Sea1-GFP reside on the vacuole membrane in wildtype and *vps501*Δ cells, however are found in the vacuole lumen (VL) in *sea1*Δ cells and *vps501*Δ*sea1*Δ cells. Note, *sea1*Δ cells appear to mask any effects of VPS501. (B) Vacuole lumen (VL) localization is defined by visually scoring the presence of GFP in the vacuole lumen. C) SEACAT subunits, Seh1-GFP and Sec13-GFP localize to the vacuole membrane in wildtype and *vps501*Δ cells, however are found enriched in non-vacuolar compartments in *sea1*Δ cells and *vps501*Δ*sea1*Δ cells. Seh1 and Sec13 have previously reported nuclear and ER roles, respectively and are likely enriched on these structures when vacuole membrane localization is compromised. (B) Vacuole lumen (VL) localization as determined by calculating the percent of cells with GFP signal on the vacuole using CMAC as a visual maker to determine vacuole boundaries. The results are from three experiments and averaged using the standard error of the mean. Indicated significance is a comparison of wildtype to single deletions or double mutants. *p<0.05, ** p<0.01, ***p<0.001 indicates significance as calculated by Student’s t-test.

**Supplemental Figure 3.**
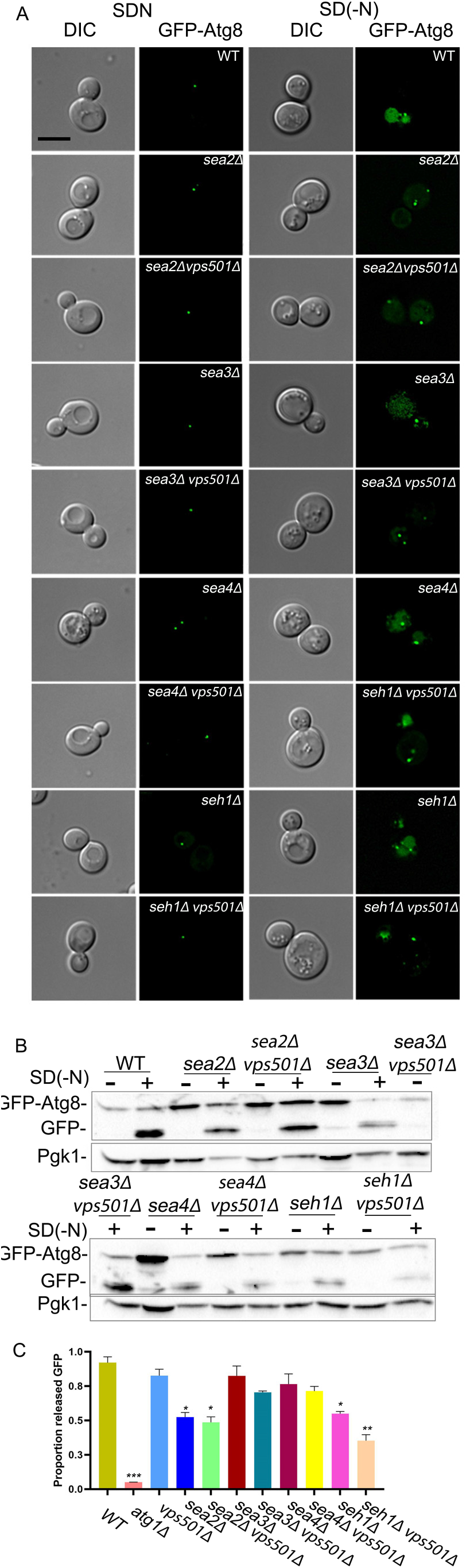
Vps501 interactions with the SEACAT complex during autophagy. (A) Maximum projection micrographs of cells expressing GFP-Atg8 in wildtype and indicated mutant cells before and after nitrogen starvation. The scale bar indicates 5μm. (B) Quantitative immunoblotting was used to detect the amount of GFP-Atg8 flux before and after autophagy induction. A partial reduction in GFP-Atg8 flux is seen when Vps501 is ablated in combination with each of the SEACAT subunits with the most significant defect occurring in *seh1Δvps501Δ* cells. A representative immunoblot is shown. Anti-Pgk1 was used as a loading control. (C) Graph of quantification of GFP-Atg8 processing. The results are from three experiments and averaged using the standard error of the mean. Indicated significance is a comparison of wildtype to single deletions or double mutants. *p<0.05, ** p<0.01, ***p<0.001 indicates significance as calculated by Student’s t-test.

